# First genetic data for the Critically Endangered Cuban endemic Zapata Rail *Cyanolimnas cerverai*, and the taxonomic implications

**DOI:** 10.1101/2022.02.07.477705

**Authors:** Alex F. Brown, Thomas J. Shannon, J. Martin Collinson, Guy M. Kirwan, Arturo Kirkconnell, Martin Stervander

**Author notes:** Correspondence (GMK) and (MS).

## Abstract

The taxonomic affinity of the near-flightless Zapata Rail *Cyanolimnas cerverai*, a Critically Endangered and highly localized species endemic to Cuba, has long been debated. Morphological analyses have suggested that this species, which constitutes a monotypic genus, could be related either to the extinct Tahitian Cave Rails (*Nesotrochis* sp.) or to the South American rail tribe Pardirallini, i.e., the genera *Neocrex, Mustelirallus*, and *Pardirallus*. Whilst pronounced phenotypic convergence–and divergence–among rails have repeatedly proven morphology-based phylogenies unreliable, thus far no attempt to sequence DNA from the enigmatic *Cyanolimnas* has succeeded. In this study, we extracted historic DNA from a museum specimen collected in 1927 and sequenced multiple short fragments that allowed us to assemble a partial sequence of the mitochondrial cytochrome oxidase I gene. Phylogenetic analyses confirm that *Cyanolimnas* belongs in tribe Pardirallini as sister to genus *Neocrex*, from which it diverged about six million years ago. Their divergence from *Mustelirallus* was estimated at about eight million years ago. Based on morphology and our mitochondrial phylogeny, we conclude that it is unjustified to retain the monotypic genus *Cyanolimnas* and tentatively recommend that *C. cerverai* and the two *Neocrex* species are ascribed to genus *Mustelirallus*.

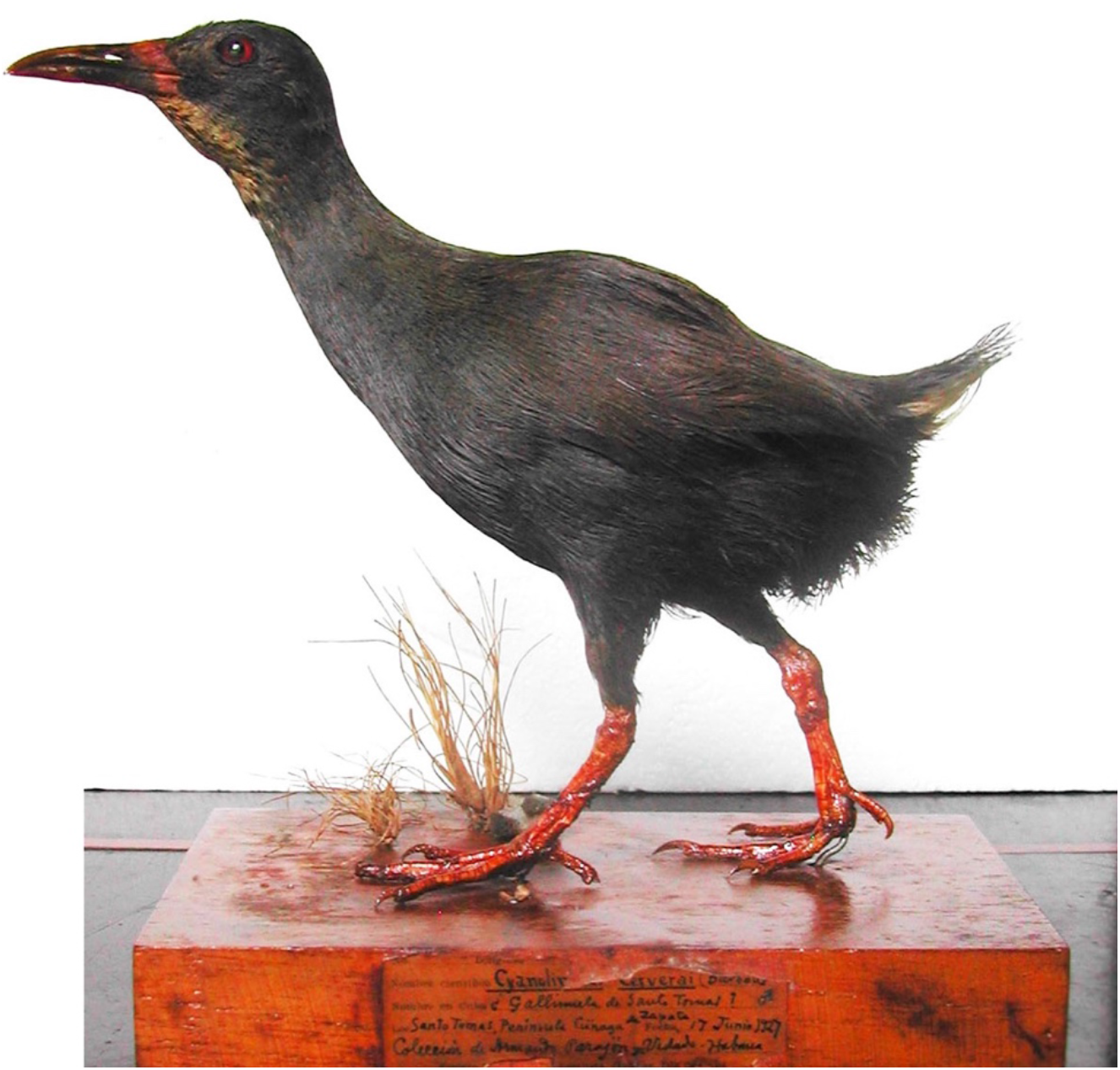

## INTRODUCTION

The Critically Endangered Zapata Rail *Cyanolimnas cerverai* is endemic to Cuba and unquestionably is one of the most poorly known birds in the West Indian region (Kirwan *et al.* 2019, Kirkconnell *et al*. 2020, Taylor *et al*. 2020). Its declining population is apparently threatened by dry-season burning of its marsh habitat, predation by introduced small Indian mongooses (*Herpestes auropunctatus*), black rats (*Rattus rattus*) and African Catfish *Clarias gariepinus* (Collar *et al*. 1992, Kirkconnell 2012, Taylor *et al*. 2020), and habitat change engendered by the spread of the invasive broad-leaved paperback (*Melaluca quinquenervia*). Nowadays, this rail is known from just six closely spaced localities in the northern Ciénaga de Zapata (Matanzas province) in south mainland Cuba, based on 14 specimens collected between 1927 and 1934 (all but one in four US museums), and a relatively small number of sight records since 1979, most recently in November 2014 (Kirkconnell *et al*. 2020). However, fossil material of *Cyanolimnas cerverai* dating from the Holocene has been identified from Cueva de Pío Domingo, Sumidero, and Cueva El Abrón, Sierra de La Güira (Pinar del Río province), Cueva de Paredones and Cueva de Sandoval (Artemisa province), Calabazar (La Habana province), Cueva de Insunsa, Cuevas de Las Charcas and Cueva del Indio (all in Mayabeque province), near Jagüey Grande (Matanzas province), Cueva de Humboldt and Cueva del Salón (Sancti Spíritus province), and the Sierra de Caballos in the northern Isle of Pines (Olson 1974, Arredondo 1984, Suárez in press), i.e., over a much larger area than the known modern range.

Genus *Cyanolimnas* Barbour & J. L. Peters, 1927, has always been considered monospecific, and was described as “A medium-sized ralline with short rounded wing; very short tail, the barbs of the rectrices very sparse; tarsus stout and short, not exceeding middle toe with claw. Bill moderate, somewhat longer than head, swollen basally.… The combination of short wing and stout tarsus suggests relationships with *Nesotrochis* Wetmore [a genus comprising three large, flightless, extinct species from the Greater Antilles]… but the latter has a tarsus more than twice as long” (Barbour & Peters 1927). Ridgway & Friedmann (1941) proffered a detailed morphological description of this genus, and thought it was apparently flightless. However, whilst *Cyanolimnas* has reduced powers of flight, it is volant (AK pers. obs.), and in any case flightlessness is now known to have arisen many times in Rallidae and cannot be used as a predictor of relationships (e.g., Olson 1973, Slikas *et al*. 2002, Kirchman 2012, Gaspar *et al*. 2020, Garcia-R & Matzke 2021, although *Cyanolimnas* was incorrectly treated as flightless in the latter). As noted by Olson (1973) and Steadman *et al*. (2013), in its robust, deep-based bill *Cyanolimnas* is similar to the two species frequently assigned to *Neocrex* Sclater & Salvin, 1868, Colombian Crake *N. colombiana* and Paint-billed Crake *N. erythrops* (both of which are now often placed in an expanded *Mustelirallus* Bonaparte, 1856; e.g., Kirchman *et al*. 2021). Plumage and osteological characters are similar to either *Neocrex*, or less so to the latter’s sister taxon *Pardirallus* Bonaparte, 1856, to which genus *Cyanolimnas* was considered most closely related in the morphological phylogenies of Livezey (1998) and Garcia-R & Matzke (2021).

Its ecology and natural history are virtually unknown; for example, a nest with eggs ascribed to *Cyanolimnas*, found in early September 1982 (Bond 1984), seems unlikely to have been identified correctly (Kirkconnell *et al*. 2020), and even this rail’s voice is unknown. A published sound recording, originally believed to pertain to *Cyanolimnas* (Reynard & Garrido 1988, Hardy *et al*. 1996), is now identified as belonging to Spotted Rail *Pardirallus maculatus*, which is a rather abundant species in the Ciénaga de Zapata (Kirkconnell *et al*. 2020). During November–December 1998 a survey using the published sound recording as a tool estimated a population of 70–90 individuals of *Cyanolimnas* (Kirkconnell *et al*. 1999). Subsequently, it was realised that the recording actually involved *P. maculatus* (Kirkconnell *et al*. 2005) and that the description of *Cyanolimnas* vocalizations in Kirkconnell *et al*. (1999) was erroneous. Inferences concerning the relationships of *Cyanolimnas* based on ecology or vocalizations are consequently impossible.

To date, just one molecular phylogenetic study, using ultra-conserved elements (UCEs), has attempted to ascertain the relationships of *Cyanolimnas* (using a toe pad from AMNH 300416, a female collected by P. Quintaña in April 1934), but was unsuccessful in yielding any UCEs, and consequently failed to place the species (Kirchman *et al*. 2021). As a result, Kirchman *et al*. (2021) proposed to treat *Cyanolimnas* as genus incertae sedis; the aim of the present work is to address this shortfall in knowledge.

## METHODS

### Primer design and evaluation

We obtained cytochrome oxidase I (COI) sequences of rallid species NCBI/GenBank (https://www.ncbi.nlm.nih.gov/nucleotide/), aligned these sequences using CLC Sequence viewer v6.0 (QIAGEN; UK) (https://digitalinsights.qiagen.com/products/clc-sequence-viewer-direct-download/) and identified regions of high similarity and roughly 50% GC content. We manually designed degenerated primers of 21–23 bp length to produce <200 bp fragments from the degraded DNA (Table 1) and tested the primer pairs on Common Moorhen *Gallinula chloropus* and Allen’s Gallinule *Porphyrio alleni* in order to preserve the DNA from the Zapata Rail.

**Table 1.**
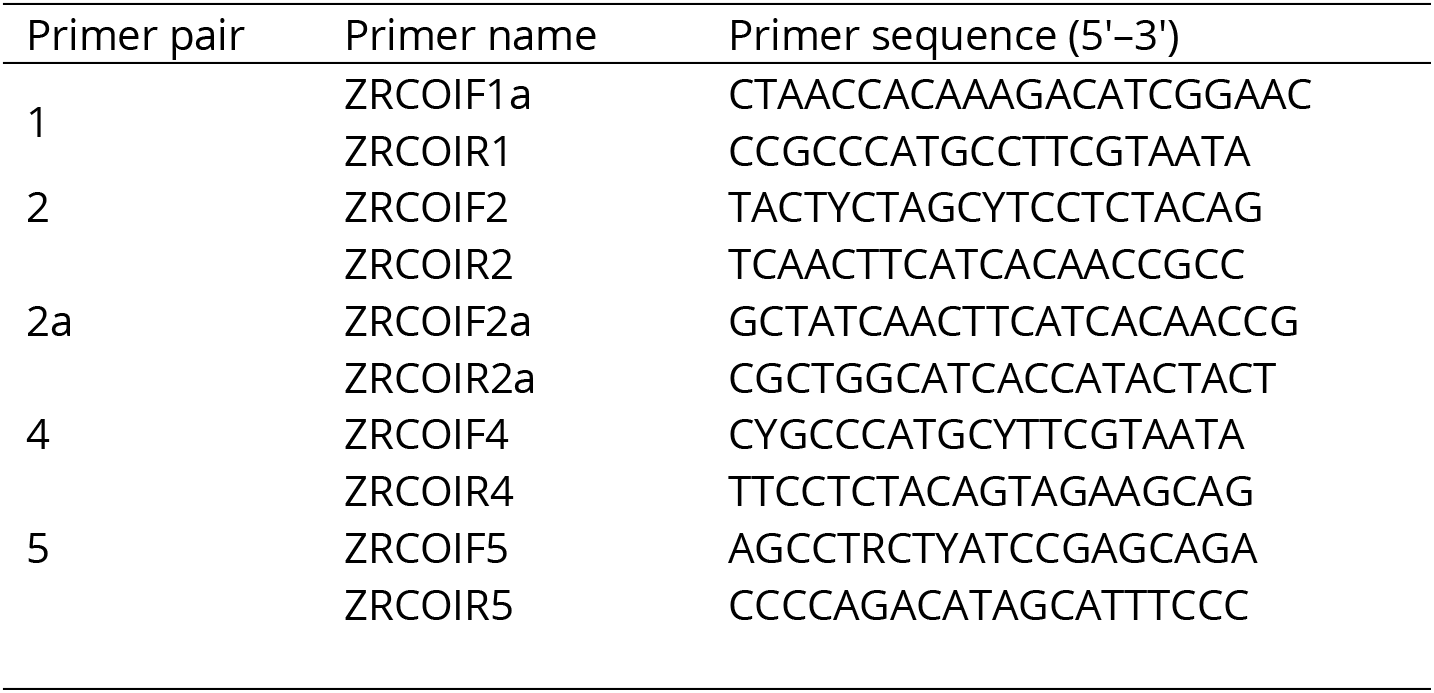
Primers designed for the Zapata Rail COI gene. This table lists all the PCR primers designed to amplify the DNA sequence of the Zapata Rail’s COI gene.

### Sampling and laboratory procedures

We obtained a toepad sample of Zapata Rail from the Museo de Historia Natural ‘Felipe Poey’, La Habana, Cuba (MFP 14.000218) and extracted DNA using QIAGEN QIAamp^®^ DNA microkit, following the manufacturer’s instructions, modified for digestion overnight at 56°C and repeated vortexing. The DNA was eluted in 80 μl Buffer AE.

We set up 50 μl PCR reactions using 2.5 μl template DNA and 47.5 μl PCR mastermix made from 5 μl of 10× OptiBuffer (Bioline Reagents Ltd, UK), 2.5 μl of primer at 10 μM, 1 μl 50 mM magnesium chloride, 0.6 μl 10 mM DNT mix (Sigma, Poole, UK), 0.5 μl BIO-X-ACT short DNA polymerase (Bioline Reagents Ltd, London, UK), 35.4 μl ddH2O. We ran samples and negative controls in PCR programs following Hebert *et al*. (2004), with a low annealing temperature in first 5 cycles to accommodate potential primer–sample mismatches: denaturing 60 s at 94°C, annealing 60 s at 45°C, and extension 50 s at 72°C. For the following 30 cycles, the annealing temperature was increased to 51°C.

Four μl 100-bp DNA ladder (Promega, Southampton, UK) and 40 μl of the PCR products were loaded with 5 μl Promega 6× DNA loading dye on 1.5% agarose gels (peqGOLD universal agarose; Peqlab, Fareham UK), stained with 1–2 μl ethidium bromide, in 1× TAE buffer. We ran the electrophoresis at 86 V, 100 mA for 20–40 minutes, inspected the gel under UV light, and cut our bands of expected size with a scalpel. The PCR product was isolated from gel fragments with a Gel Extraction Kit (Qiagen, Manchester, UK), following the manufacturer’s instructions, and eluted in 30 μl elution buffer EB. We quantified the purified PCR product with a Nanodrop spectrophotometer and diluted the PCR products to 10 μg/μl, after which they were bidirectionally Sanger sequenced by Source Bioscience (Nottingham, UK).

### Phylogenetic analyses

We initially scrutinized the sequences using FinchTV (Geospiza Inc.) and then assembled them to the rallid sequence set used for primer design in CLC Sequence Viewer 8 (https://www.qiagenbioinformatics.com/products/clc-sequence-viewer/), followed by extraction of the 601-bp long overlapping Zapata Rail COI consensus sequence. We then obtained a larger set of 450 COI sequences of extant taxa within the order Gruiformes from GenBank, and additionally included sequence data from ancient DNA of the extinct *Nesotrochis steganinos* (Oswald *et al*. 2020), because of its proposed close relationship to *Cyanolimnas* in combination with its recent extinction (Oswald *et al*. 2020 dated bones to 6,430 ± 30 years before present). We aligned sequences with MAFFT (Katoh & Standley 2013) in Geneious v. 10.2.6 (Kearse *et al*. 2012) and selected ≤3 per taxon based on length and quality (for taxa and accession numbers, see Table S1). We further trimmed this dataset to a 962-bp region that maximized overall sequence coverage and fully overlapped the Zapata Rail COI sequence (see Data Availability). We employed the greedy algorithm (Lanfear *et al*. 2012) in PartitionFinder2 (Lanfear *et al*. 2016) to evaluate the full set of SNA substitution models and explore optimal partitioning with regard to codon position, using PhyML (Guindon *et al*. 2010). The best partitioning scheme by AICc was identified as individual substitution models per codon position: the general time-reversible (GTR; Tavaré 1986) model with the among-site rate variation following a gamma (Γ) distribution and with a proportion of invariant sites (I) for position 1, the transversion model (TVM) + Γ + I for position 2, and the transition model (TiM) + Γ + I for position 3.

We implemented the partitioning scheme in Beast v. 2.6.6 (Bouckaert *et al*. 2014) using the plugin SSM v. 1.1.0 and set the proportion of invariant sites to be estimated, and the Γ distribution to be estimated across four rate categories. Employing a birth–death speciation model, we set the prior for death rate to follow an exponential distribution with mean 1. We enforced monophyly of the clades Rallidae, Ralloidea, and Gruoidea, and employed dating calibration priors on Gruiformes and stem Rallidae following Chaves *et al*. (2020). We then ran two iterations of 10 million generations, sampling every 1,000 generations, and optimized tuning parameters and operator weights to produce the final xml specification used for our analyses (see Data Availability). We ran five replicate analyses at different seeds in Beast, sampling every 1,000 generation for 20-60 million generations, until inspection in Tracer v. 1.7.1 (Rambaut *et al*. 2018) revealed stationarity and sufficient effective samples sizes (all parameters >200). Some substitution model-related parameters for codon position 1 and to some degree 2 (specific substitution rates, gamma shape, and proportion invariant sites) seemed to have dual optima, with shifting within and between replicates. Thus, replicates converged in either of two groups; however, this had no impact on the topology, support, or dating of the focal clade. We calculated maximum clade credibility trees with mean node heights after discarding 5–50% as burn-in.

In addition to the Bayesian inference (BI) with Beast, we also used IQtree v. 2.1.2 (Lanfear *et al*. 2020) for phylogenetic analyses based on maximum likelihood (ML). We applied the partitioning and substitution model scheme determined with PartitionFinder2 and ran 1,000 ultrafast bootstrap replicates.

## RESULTS

All phylogenetic analyses placed *Cyanolimnas* as sister to *N. erythrops*, the pair of which was sister to the Ash-throated Crake *Mustelirallus* (formerly *Porzana*) *albicollis* (Figure 1). These three taxa formed a sister clade to the genus *Pardirallus* (Figure 1). There was near-full support for a *Pardirallus–Mustelirallus–Neocrex–Cyanolimnas* clade (posterior probability (PP) 1.0 in the BI analyses and 99% ultrafast bootstrap support (UFB) in the ML analyses). The exact placement of *Cyanolimnas* as sister to *Neocrex* received full support (UFB 100%) in the ML analyses and PP 0.86 with BI (Figure 1). There were some differences in inferred topology between BI replicates (see Data Availability); however, none of these occurred within the focal *Pardirallus– Mustelirallus–Neocrex–Cyanolimnas* clade.

**Figure 1.**
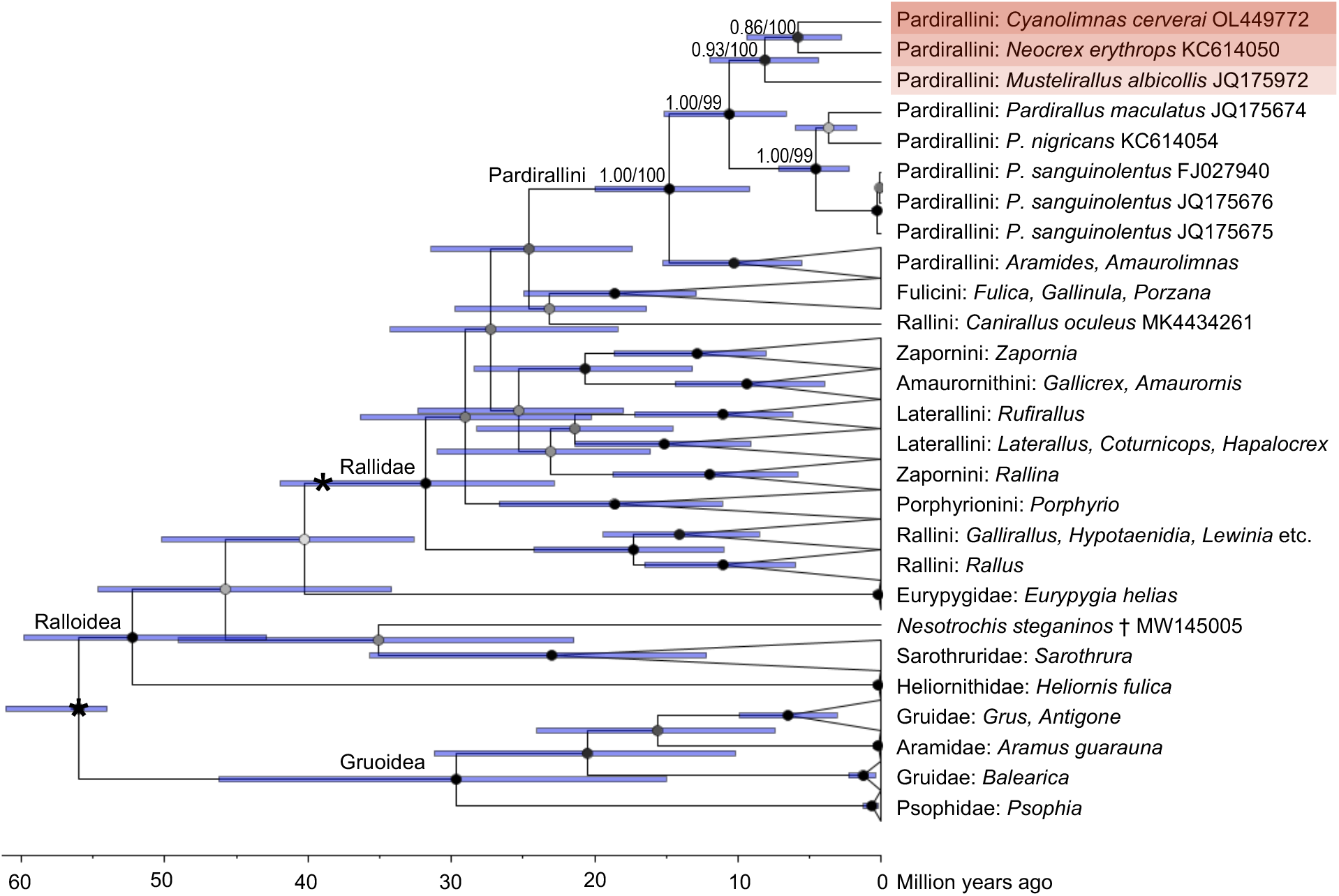
Phylogenetic tree of Gruiformes, based on mitochondrial cytochrome oxidase I (COI) sequences, analyzed with Bayesian inference (BI) in Beast. Posterior probability is indicated by node colour (0.0 = white; 1.0 = black) and specified with labels at nodes for the focal clade in tribe Pardirallini (first number), together with ultrafast bootstrap support values (0–100) from maximum likelihood (ML) analyses with IQtree (second number). The asterisks at the root of Gruiformes and the stem of Rallidae indicate fossil calibration points. Non-focal clades represented by multiple sequences have been collapsed and labelled according to Kirchman et al. (2021) with tribe (for Rallidae) or family (outside Rallidae) followed by species or genus (if multiple species). For non-collapsed replicate BI trees and ML tree, see Data Availability.

The inferred mean age of split between *Cyanolimnas* and *Neocrex* was 5.9 MA (95% highest posterior density 2.8–9.4 MA), between those and *Mustelirallus* 8.1 (4.4–12.0) MA, and between that clade and *Pardirallus* 10.6 (6.6–15.2) MA (Figure 1).

*Nesotrochis* was recovered as a sister lineage to Sarothruridae (Figure 1).

## DISCUSSION

We provide the first genetic data for *Cyanolimnas*, as the species has not been included or successfully sequenced by any previous molecular-phylogenetic study of the Rallidae (e.g., Garcia-R *et al*. 2014, Garcia-R & Matzke 2021, Kirchman *et al*. 2021), which led Kirchman *et al*. (2021) to consider its placement ‘incertae sedis’. It should be stressed that the reliability of phylogenies based on single genetic markers is limited, and this is particularly true for mitochondrial markers. The fast-evolving mitochondrion occurs with multiple copies per cell and has historically been widely used for molecular phylogenies, and further remains important for genetic barcoding. However, for several reasons the mitochondrial phylogenetic signal may be discordant from that of the nuclear genome (e.g., through introgression) and reflect an evolutionary trajectory different from ‘true’ speciation events (Toews & Brelsford 2012). In our analyses, for example, the COI tree renders Gruidae paraphyletic with respect to Aramidae (Figure 1). Nevertheless, mitochondrial data are relevant for a first test of presumed phylogenetic context, and yield useful relative divergence times.

Similar to the whole-mitochondrion analyses by Oswald *et al*. (2020), our analyses of COI placed *Nesotrochis* outside Rallidae, separated from *Cyanolimnas* by some 45 million years (Figure 1). Our data instead provide strong support for the previous hypotheses rooted in morphological traits that have placed *Cyanolimnas* in what has been referred to as the *Aramides* clade’ (Livezey 1998, Garcia-R *et al*. 2014, Garcia-R & Matzke 2021), better termed tribe Pardirallini *sensu* Kirchman *et al*. (2021). Within this grouping, the species appears to be most closely related to *N. erythrops* and *M. albicollis*, but less so to the three species of *Pardirallus*, in partial congruence with prior predictions (Olson 1973, Livezey 1998, Garcia-R & Matzke 2021). Although it is now rather well established that plumage is not necessarily a reliable character for inferring phylogenetic relationships within the Rallidae (e.g., Garcia-R *et al*. 2014, Stervander *et al*. 2019, Chaves *et al*. 2020), in the present case morphology appears to be rather informative.

The key question that arises is whether the separate monospecific genus *Cyanolimnas* is warranted? The age of the split between *Cyanolimnas* and *Neocrex* (5.9 MA) is comparatively young relative to many other Rallidae, a family that contains several old generic clades (e.g., *Aramides* 7.4 MA, *Porzana* 9.8 MA, *Rallus* 11.0 MA, *Rallina* 12.0 MA, *Zapornia* 12.9 MA, and *Porphyrio* 18.6 MA based our COI analyses; Fig. 1). However, some clades contain even younger taxa, for example the *Eulabeornis–Cabalus–Gallirallus–Hypotaenidia* group of the Rallini (which arose 3.4-4.1 MA), all or most of which genera are accepted by some authorities (e.g., Dickinson & Remsen 2013, del Hoyo & Collar 2014, Kirchman *et al*. 2021). Nevertheless, in the latter case these four genera are sometimes treated alternatively as a single genus, for which the name *Gallirallus* Lafresnaye, 1841, has priority (Kirchman 2012).

We consider that the available molecular and morphological evidence in combination strongly supports that *C. cerverai* be included in either genus *Neocrex* Sclater & Salvin, 1869, or *Mustelirallus*, Bonaparte, 1856, given that its most divergent trait from either of these two, (near-)flightlessness, is well accepted to be not taxonomically informative (Olson 1973, Slikas *et al*. 2002, Kirchman 2012, Gaspar *et al*. 2020, Garcia-R & Matzke 2021). Although *N. colombiana*, an exceptionally poorly known rallid of northwestern South America and easternmost Panama, has yet to be sampled genetically, to date there is no indication that it is not very closely related to *N. erythrops* (Wetmore 1967, Olson 1973, Taylor 1996). Following the lead of Kirchman *et al*. (2021), we recommend that, for the present, the most parsimonious approach is to treat all four species—Ash-throated Crake, Paint-billed Crake, Colombian Crake and, now, Zapata Rail—as members of a single genus, for which the name *Mustelirallus* (masculine) has priority. The new combinations are *Mustelirallus albicollis, M. erythrops, M. colombianus*, and *M. cerverai*. Kirchman *et al*. (2021) incorrectly listed Colombian Crake as *M. col<u>u</u>mbianus*, overlooking that Bangs’ (1898) original spelling was *Neocrex col<u>o</u>mbianus* as noted by Dickinson & Remsen (2013).

## Acknowledgements

GMK and AK are grateful to staff, particularly Ianela García-Lau, Manolo Barro and volunteers at the Museo de Historia Natural ‘Felipe Poey’, La Habana, Cuba, for access to relevant specimens. This study was funded by University of Aberdeen (AB) and The Sound Approach Ph.D. Studentship (TJS).

## Author contributions

Conceived and managed project: GMK, JMC, MS. Provided biological sample: AK, GMK. Designed primers, performed molecular lab work: AFB, TJS, JMC. Assembled COI sequence: AFB, TJS. Performed phylogenetic analyses: MS. Wrote the manuscript: GMK, MS.

## DATA AVAILABILITY

The Zapata Rail COI sequence has been deposited at Genbank with accession number OL449772. Supporting material has been deposited at Zenodo and is available at https://doi.org/10.5281/zenodo.5805456. This includes Table S1 (taxon name, classification, and accession numbers for sequences included); sequence alignment nexus file; Beast xml input file; Beast output including maximum clade credibility tree files; log files and raw tree files; and IQtree consensus tree file.

